# High permeability of the CSF flow in the juvenile mouse brain by brain-wide DCE-MRI

**DOI:** 10.1101/2021.09.21.461189

**Authors:** Yayoi Wada, Hirohiko Imai, Yuki Yamawaki, Md Sorwer Alam Parvez, Sae Matsui, Gen Ohtsuki

## Abstract

The vasculature system with a lymphatic function in the brain is manifested as meningeal lymphatic vessels and the glymphatic system, which drain waste products from cerebrospinal fluid (CSF) and produce interstitial fluid. Invasion of stimulated immune cells or inflammatory cytokines during the maturation is regarded as a sign of the disruption of brain functions through excessive immune stresses. However, it is unclear when the lymphatic system is functionally completed and in which parts of the organ the brain immunity is privileged (*i.e*., the blood-brain barrier is fully established). Here, we visualized the whole-head signal of gadolinium (Gd) contrast agents in mouse brains to investigate the CSF flow from the signals of Gd-contrast agents using magnetic resonance imaging (MRI) in the course of developmental stages. We found higher Gd-signals in the olfactory bulbs, prefrontal cortex, parietal surface regions of the neocortex along with dorsal meningeal lymphatic vessels, ventral midbrain, and a part of cerebellum, as well as in the basal brain regions, at the immature stage of postnatal (P) 4 weeks, compared to the P8-12 weeks. Our results suggest that the barrier of the vasculature system in mouse brains is still permeable until P1-month and small molecules can leak into the parenchyma.

## Introduction

The blood-brain barrier (BBB) is a sophisticated endothelial barrier that controls the milieu of the brain and prevents the extravasation of fluids, solutes, inflammatory cytokines, microbial pathogens, blood- and immune-cells from the blood flow, as an immune privilege. At the BBB, the water component of cerebrospinal fluid (CSF) dominantly crosses the astrocytic water channels AQP-4 (Nielsen et al., 1997), and the solutes enter the brain parenchyma via astrocytic transporters or ion channels. At the paravascular space between the basement membrane of smooth muscle cells and glia limitans, the CSF infiltrates into the brain parenchyma as the interstitial fluid (ISF), which drives the removal of waste products, gasses, and extracellular solutes of the parenchyma (Iliff et al., 2012, 2013; Louveau et al., 2017). According to Kiviniemi et al. (2016), cardiac pulsation (~1.1 Hz), respiratory pulsation (~0.4 Hz), and very low-frequency vasomotor pulsations (< 0.01 Hz) (Kiviniemi et al., 2016) drive CSF/ISF exchanges. The BBB is formed during embryogenesis in the murine but might not be completely impermeable to solutes and cells during the postnatal period. After birth, the junction of BBB matures very rapidly, and they are thought to attain their adult characteristics (Bauer H et al., 1995; Bauer HC et al., 2014). However, it is unclear when the permeability barrier is functionally completed, how CSF leakage to the parenchyma is regulated in the course of brain maturation, and if there are differences of the permeability depending on brain regions.

A principal function of CSF is to provide a constant fluid flow to maintain homeostasis in the ventricular system and on the surface of the brain by carrying lots of secreted mediators, solutes, and immune cells and excreting the waste products of the parenchyma. The CSF is also able to transmit hormones, transmitters, inflammatory cytokines, and even exosomes (Brown et al., 2004; Praetorius, 2007; Yagi et al., 2017; Lepennetier et al., 2019). The CSF is produced by the ependymal cells in the choroid plexus of all the ventricles: lateral, third, and fourth ventricles. The entire CSF volume and production rate had been estimated at 35 μl and is 0.32 μL/min, respectively, in the mouse brain (Pardridge, 2016). While CSF is produced in the choroid plexus with a rich capillary bed, the osmotic pressure of the CSF is normally around 5 mOsm/kg higher than that of the serum and the blood plasma (Cserr, 1971; Brown et al., 2004; Praetorius, 2007). According to Akaishi et al. (2020), the CSF osmolarity is 289.5 ± 6.6 mOsm/kg while the serum osmolarity is 283.6 ± 6.6 mOsm/kg, implying that the passive osmotic pressure contributes to the CSF flow, as well as the pulsatile does. A recent direct measurement with less leakage of the CSF production from lateral ventricles indicated lower production volume of 108.0 ± 7.6 nL/min and 83.9 ± 2.8 nL/min in 2-month-old male and female C57BL/6 mice, respectively (Liu et al., 2020). Previous research by Kress et al. (2014) showed that the efficiency of exchange between the subarachnoid CSF and the brain parenchyma is higher at least in young 2-month-old mice than in the aged group. The study demonstrated the decline of the CSF/ISF exchange in the course of maturation using in vivo and ex vivo fluorescence microscopy. Fluorescent CSF tracers were slowly infused into the subarachnoid CSF of the cisterna magna, and their results of tracer influx indicated the region-dependent difference in the infiltration of fluorescent CSF tracers and higher amount of infiltration in the basal brain regions of the 2-3-month-old C57BL/6 mice than 18-month-olds (Kress et al., 2014). These results suggested the idea that CSF-infiltration into brain parenchyma is higher in the immature brain and distinct brain regions are more permeable. However, the study has focused on the CSF/ISF exchanges in aged disease brains and was not intended to find the CSF leakage through brain regions thoroughly. Therefore, it remains unclear if such permeable regions are broader in younger brains and along what vasculatures the CSF infiltration is distinct. In the present study, we aimed to reveal regions with higher permeability of vasculature system in the earlier age of experimental animals with a whole-brain visualization by dynamic contrast-enhanced magnetic resonance imaging (DCE-MRI). We investigate whether Gd-signals are fairly constrained in the pathways of CSF at developmental stages by injection from the cisterna magna.

## Results

### In vivo visualization of CSF flow using DCE-MRI from mouse brains

To investigate the *in vivo* CSF flow and CSF-permeation to the parenchyma in the course of maturation in the entire vasculature system of the head, we visualized the flow dynamics and infiltration of the Gd-signal in the brain parenchyma using 7T MRI system (**Figure 1**) (**Materials & Methods**). We used a Gd-based contrast agent GadoSpin M (Miltenyi Biotec, USA), with a low molecular weight (938 g/mol), which allows visualization of the leak of CSF (Aydin et al., 2008). Upon injection into the cisterna magna (*i.e*., the posterior cerebellomedullary cistern), GadoSpin M rapidly diffuses in the space of ventricles. To investigate the leakage of Gd-signal in the entire vasculature system of the brain, we intentionally injected a high dose of Gd-contrast agent of 35−38 μl volume to fulfill the entire space of ventricles. Due to an opening called median aperture (foramen of Magendie) between the cisterna magna and the fourth ventricle, our injection into the cisterna magna made a substantial backflow to the fourth ventricle. The Gd-contrast agent reached the lateral ventricles, which was seen in the previous experiment by Da Mesquita et al. (2018a) and Ahn et al. (2019) (**Figures 1B & 1C**). Gd-signals indicate the signals in infused ventricles, para-sinus, lymphatic flow, and even parenchyma (**Figures 1 & 2**). Additionally, from the lateral aperture (foramen of Luschka) of the fourth ventricle Gd-contrast agent would have infused into the subarachnoid space. Because of the very rapid transfer of the systemic circulation by pulsations, Gd-contrast agent can circulate into the para-arteries and their signals were also visible within several ten seconds.

**Figure 1.**
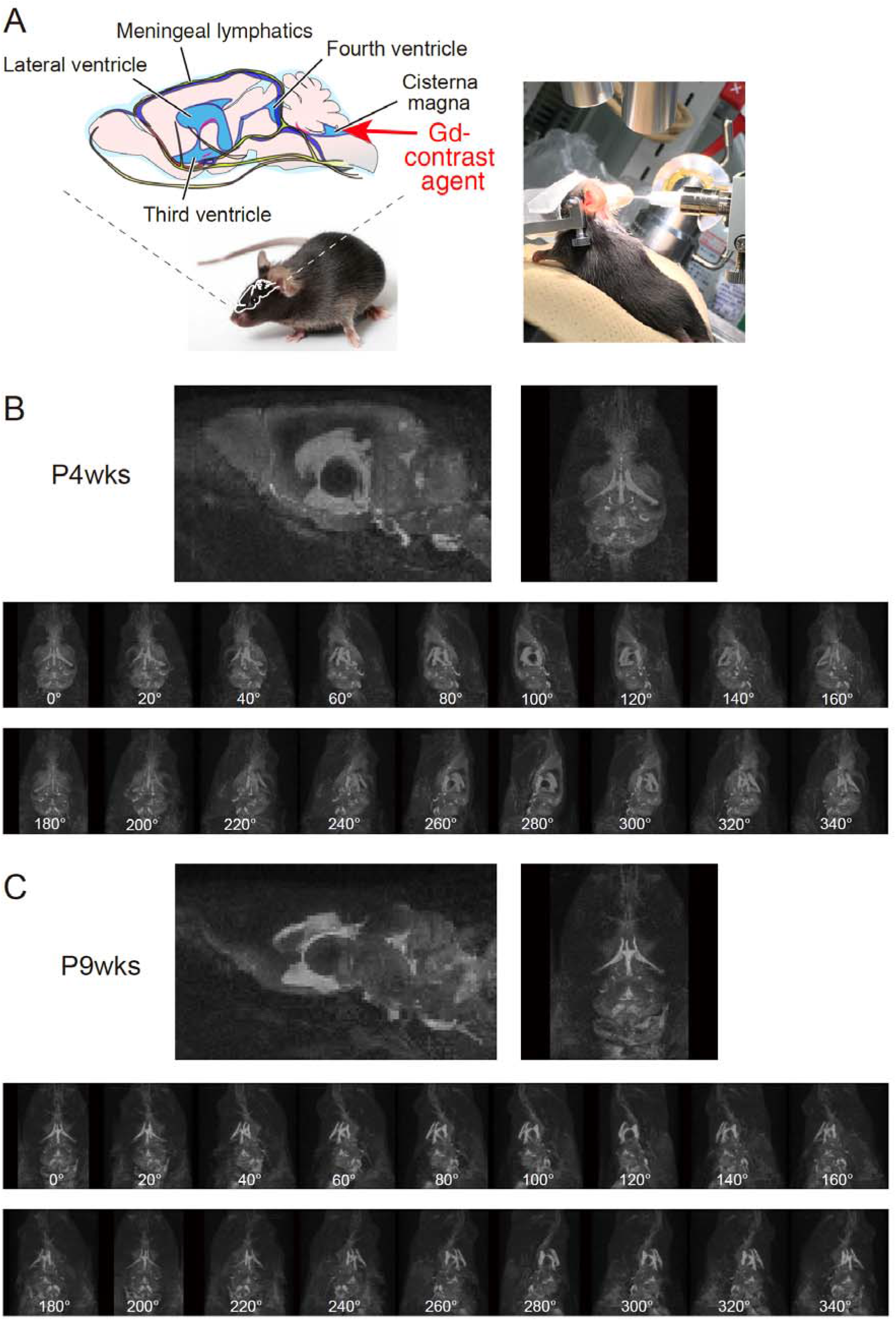
Schematic illustration of vasculature system and 3D maximum intensity projection (MIP) images of whole head T1-weighted images of young (P4-weeks) and mature (P9-weeks) C57BL/6 mice. **A,** *left*, Brain lymphatic vessels (olive green), veins & venous sinus (blue), and ventricles (sky blue) are drawn. *Right*, Operation for the injection of Gd-contrast agent. A 27-gage needle is inserted into the cisterna magna at a slow angle of negative 5 degrees from the front. After injection, we started the MR imaging within 7-9 minutes. **B, C,** Sagittal, and horizontal views of T1-weighted MIP images of the P4- and P9-week-old mice are shown (B and C, respectively). Each bottom low shows the rotation of both representative brains at 20 degrees per frame. Please note the difference of Gd-signals between P4- and P9-week-old mice. The images were captured at 9’00” (Frame 2).

**Figure 2.**
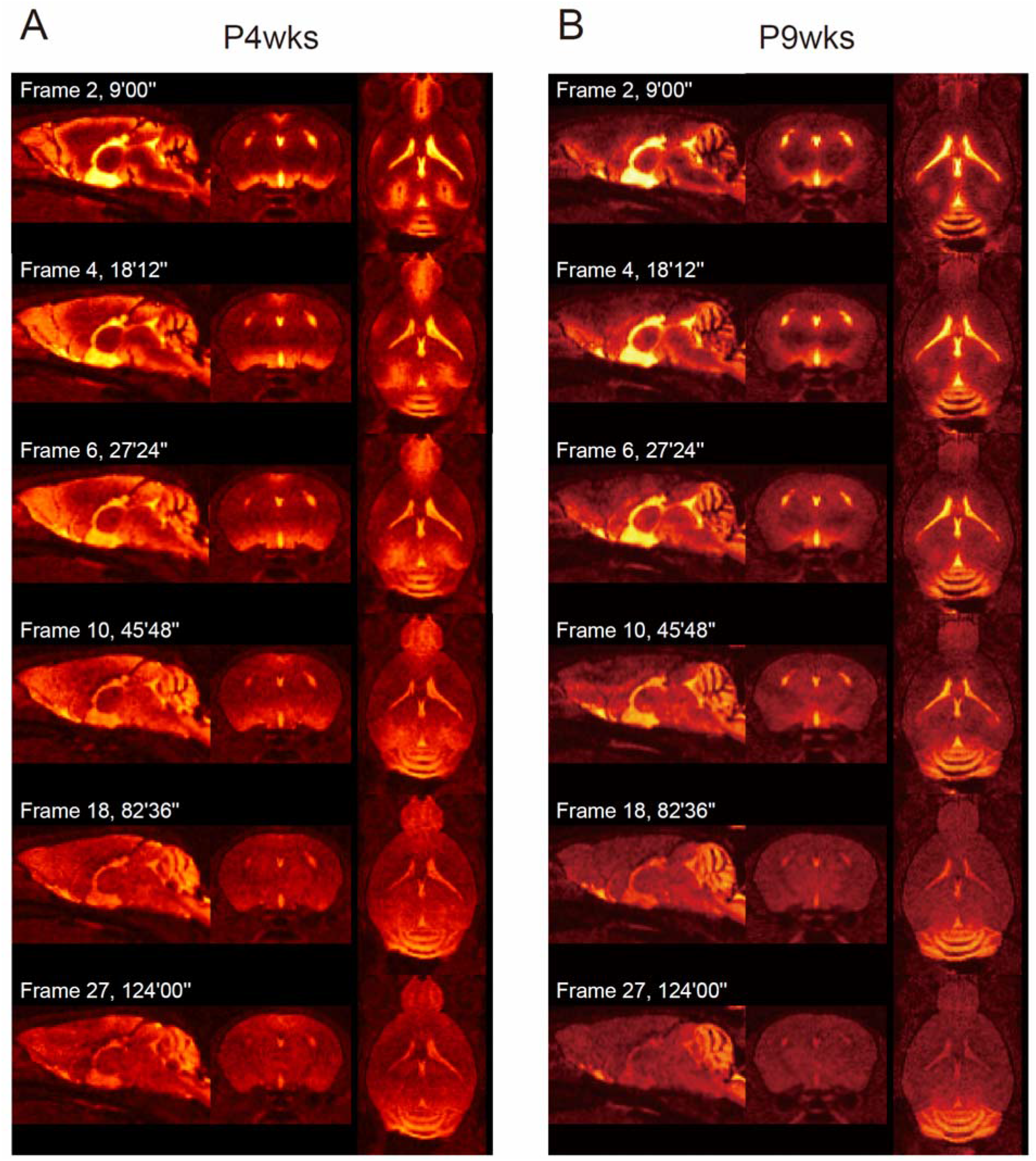
Time-lapse images of Gd-signal-spread of whole head T1-weighted images of P4- and P9-week-old mice. **A,** Representative time-lapse images of Gd-signals of the P4-week-old animal are shown at the three 2D planes. **B,** Representative time-lapse images of Gd-signals of the P9-week-old animal are shown. As a baseline image, a non-contrast-enhanced image was taken before the injection of the Gd-contrast agent. The maximum duration for the start of the initial MRI acquisition as Frame 2 was 9 minutes throughout the study, and then, we use the time in the time display.

To compare the difference in the Gd-signals in the course of maturation, we investigated signals at postnatal (P)4-weeks and P8-12 weeks (**Figures 1B & 1C**). Right after the injection, we started monitoring the Gd-signals for more than 2 hours. Gd-signals indicate a wider spread of the contrast agent in the younger P4-week brain particularly from para-sinus and lymphatic flow (**Figures 1B & 1C, Figure 2**). In the mature P8-12-week brain, the CSF bulk flow with the Gd-contrast agent injected into the cisterna magna is along the paths of the basal cistern to the olfactory artery under isoflurane anesthesia, which represents the similar observation of Stanton et al. (2021). According to Stanton et al. (2021), the Gd-contrast agent is distributed along the perivascular space of the arteries in the subarachnoid space cisterns. From there, the contrast agent either infiltrates into the parenchyma dorsally from the basal vasculature or continues as CSF bulk flow to efflux via the nasal turbinates, and the pharyngeal lymphatic vessels. Distinct from the mature mouse, the CSF flow is detected in the subarachnoid space cistern of the dorsal cerebrum in the P4-week brain (**Figures 1B & 2A**).

As shown in the T1-weighted images (: sagittal, horizontal, and rotation images of the initial flame), there are prominent differences between two developmental periods in the illuminated regions in the brains: especially, the signals in the surface of the prefrontal cortex including anterior cingulate cortex to the retrosplenial cortex along the rostral rhinal vein and the superior sagittal sinus (SSS), midbrain along the aqueduct, and olfactory bulbs (**Figures 1B & 1C**). At the parietal surface regions of the neocortex, including areas along with the rostral rhinal vein and SSS, there are the dorsal meningeal lymphatic vessels (Louveau et al., 2015). The previous studies suggested high permeability of the fluorescent molecules to parenchyma in 2-month-old animals (Perry et al., 1997; Kress et al., 2014), but these results suggest further permeability at younger one-month-old.

In Figure 2, the time-lapse images of the three 2D-planes (: Frame 2, 4, 6, 10, 18, and 27 at 9’00”, 18’12”, 27’24”, 45’48”, 82’36”, and 124’00”, respectively) show that CSF runs mainly into the basal lymphatic outflow in both ages. To measure the time courses of the Gd-signals in the whole brain regions, we first monitored the signals along the cisternae, ventricles, and sinuses (**Figure 3**). We took 30 points of interest of those lymphatic vessels and ventricles (**Extended Data Figure 3-1**), and we measured the Gd-signals along the pathway: cistern magna, aqueduct, the third ventricle, lateral ventricle, olfactory artery, SSS, and other vasculatures. Obtained time courses of Gd-signals in each region were normalized with the value of the maximum intensity of all the regions and mean ± SEM are shown in **Figure 3A** as “Lymph & Ventricles”.

**Figure 3.**
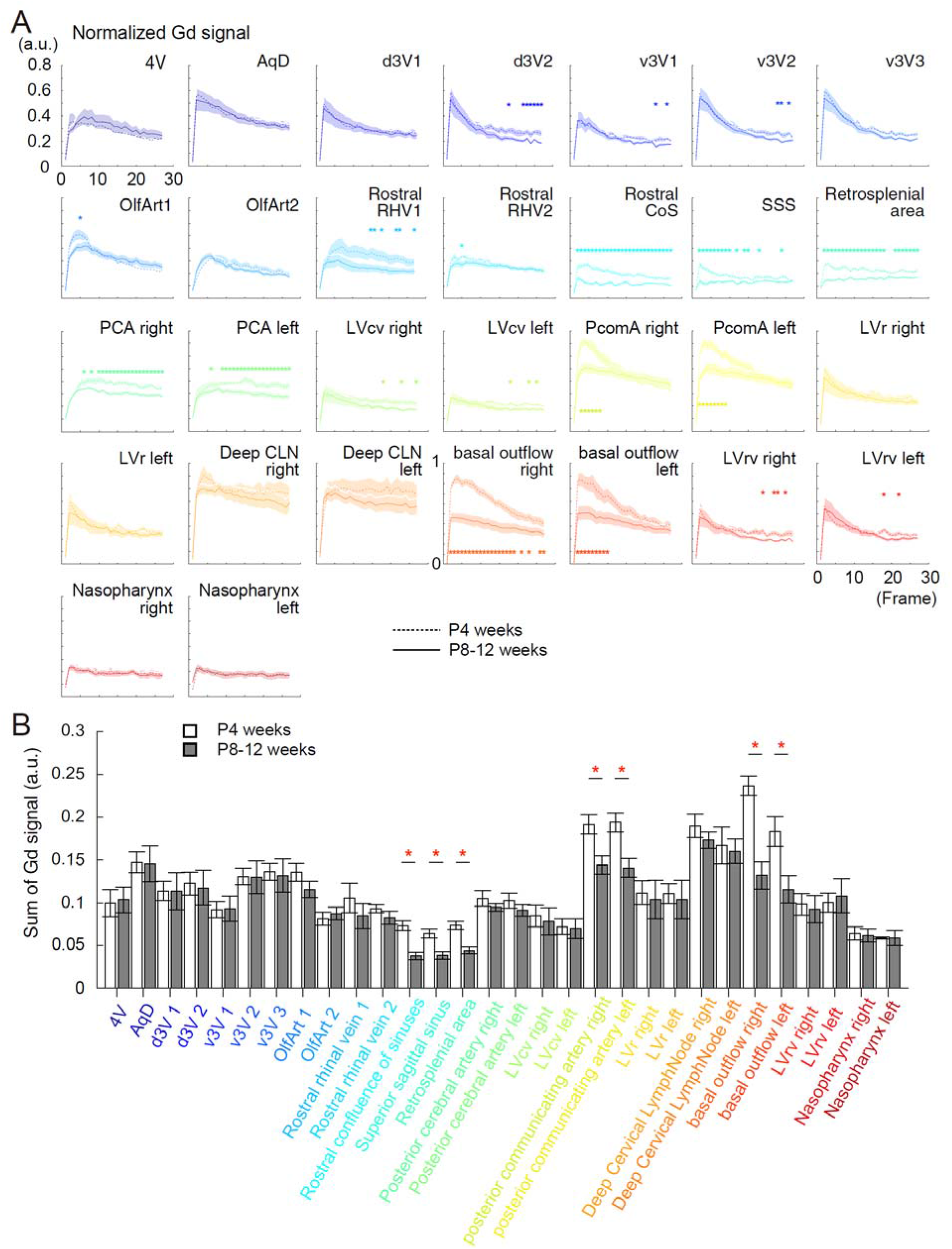
Increases in the Gd-signal of immature brains in the distinct regions along to the vasculature systems. From the lymphatic vessels or ventricles (“Lymph & Ventricles”), we took 30 points of interest (**Extended Data Figure 3-1**). **A,** Time courses of the normalized Gd signals at different brain regions of the lymphatic vessels and ventricles at two developmental stages (P4 and P8-12-weeks). Frame 1 is taken before injecting the Gd-contrast agent. Frames 2-27 were taken with an interval of 4 min 36 s. The time courses of Gd-signals are shown as mean ± SEM (mean: dotted line, P4-weeks; solid line, P8-12-weeks, ± SEM, delineated shadow). The time points with a significant difference between the P4- and P8-12-weeks are asterisked (*p < 0.05). **B,** Summary bar graphs of accumulated Gd-signals for 36 min 48 s (from Frame 2 to Frame 9) at two developmental stages of each region (mean ± SEM, blank and filled bars for P4- and P8-12-weeks, respectively). Significant differences of “Lymph & Ventricles” were found in the Rostral CoS (ranksum = 70, *p = 0.001), SSS (ranksum = 75, *p = 0.0043), retrosplenial area (ranksum = 71, *p = 0.0014), PcomA Right (ranksum = 78, *p = 0.0093), PcomA Left (ranksum = 76, *p = 0.0056), basal outflow Right (ranksum = 69, *p = 0.0008), and basal outflow Left (ranksum = 82, *p = 0.0282). Otherwise, not significant. 4V, fourth ventricle. AqD, aqueduct. d3V, dorsal third ventricle. v3V, ventral third ventricle. OlfArt, olfactory artery. Rostral RHV, rostral rhinal vein. Rostral CoS, rostral confluence of sinuses. SSS, superior sagittal sinus. PCA, posterior cerebral artery. PcomA, posterior communicating artery. LVcv, caudal ventral lateral ventricle. LVrv, rostral ventral lateral ventricle. LVr, rostral lateral ventricle. dCLN, deep cervical lymph node.

**Extended Data Figure 3-1.**
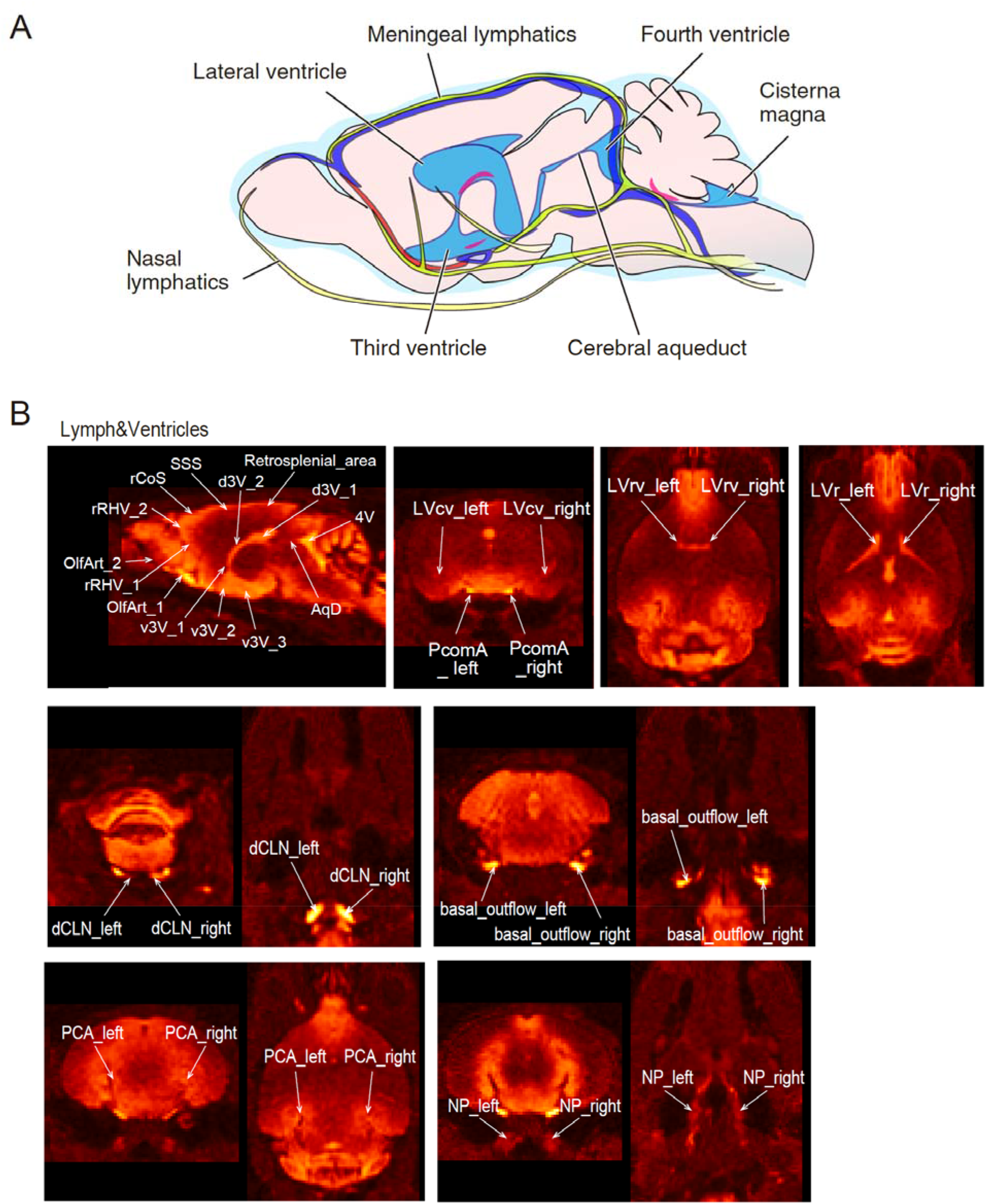
Schematic illustration of vasculature system and illustrations of brain lymphatic vessels and ventricles with interest, Related to Figure 3. 4V, fourth ventricle. d3V, dorsal third ventricle. v3V, ventral third ventricle. AqD, aqueduct. dCLN, deep cervical lymph node. rCoS, rostral confluence of sinuses. LVcv, caudal ventral lateral ventricle. LVrv, rostral ventral lateral ventricle. LVr, rostral lateral ventricle. NP, nasopharynx. OlfArt, olfactory artery. PCA, posterior cerebral artery. PcomA, posterior communicating artery. rRHV, rostral rhinal vein. SSS, superior sagittal sinus.

There is a wide range of variance in the normalized Gs-signals across the regions of lymphatic vessels & ventricles in each developmental stage. The normalized Gd-signals were high in the basal brain regions and low in the dorsal brain regions. While the deep cervical lymphatic nodes showed a rapid increase in the signals after Gd-injection, the signals remained high for a long duration, representing the convergence of the brain drainage system of the lymphatic nodes. The posterior communicating artery (PcomA) close to the pituitary recesses and the basal outflow also showed high Gd-signals initially, but the time course of signals suggests a gradual attenuation of the drainage (**Figure 3A**). We found the prominent difference between P4-week- and P8-12-week-old mice in the signals of the basal outflows and PcomA, implying faster excretion of the brain CSF in the young animals. In contrast, the other gland signals of the deep cervical lymphatic nodes and nasopharynx did not show any significant differences between both developmental periods (**Figure 3**). In Figure 3B, the bar graphs of the cumulative normalized Gd-signals of the initial 8 Frames (: from Frame 2 to Frame 9, for 36 min 48 s), again, indicate the distinct regions with high Gd-signals in the young period. Along the regions of lymphatic vessels & ventricles, there are the substantial difference with significance between P4-week- and P8-12-week-old mice in the time points of Gd-signals of the dorsal and ventral third ventricles, olfactory arteries, rostral rhinal veins, rostral confluence of sinuses (CoS), SSS, retrosplenial area, posterior cerebral arteries, and caudal ventral lateral ventricle. Notably, P4-week mice showed higher intensity in the signals of the rostral CoS, SSS, and retrosplenial area than those of P-8-12 week mice (**Figure 3B**). Comparison of the time to peak (**Figure 5A**; **Extended Data Figures 5-1A&B**) and FWHM (**Figure 5B**; **Extended Data Figure 5-1A&B**) of Gd-signals also suggests the higher extent of Gd-signals in rostral rhinal vein, rostral CoS, SSS, and retrosplenial area in P4-week group. These results suggest that the regions along the basal artery to the rostral vein and retrosplenial areas have leaky ducts at P4-weeks, whereas the leakage is shut down at P8-to P12-weeks.

**Figure 4.**
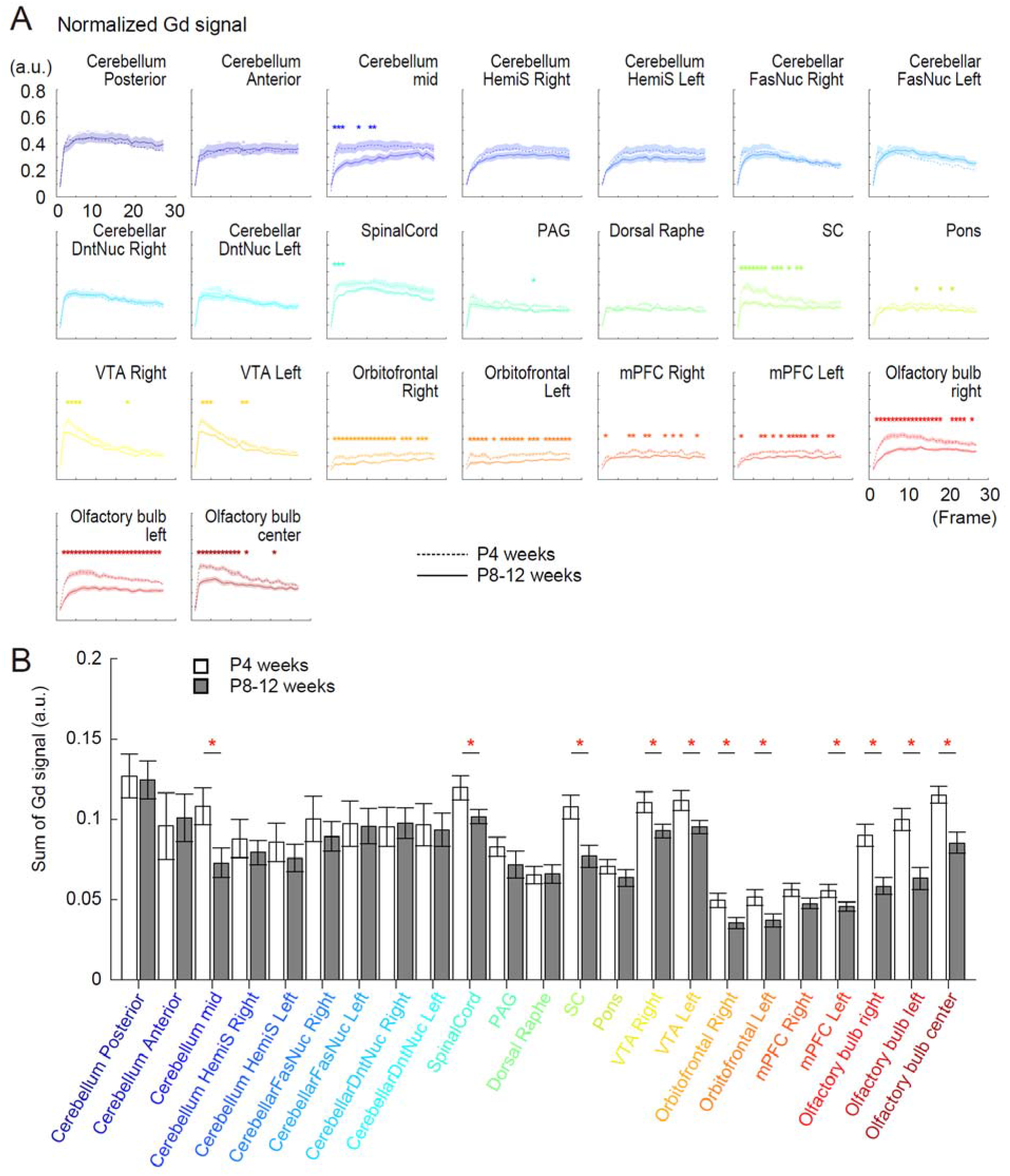
Increases in the Gd-signal of immature brains in the distinct regions of the brain parenchymas. From the brain parenchymas (“Parenchyma”), we took 23 points of interest (**Extended Data Figure 4-1**). **A,** Time courses of the normalized Gd signals at different brain regions of parenchymas at two developmental stages (P4 and P8-12-weeks). Frame 1 is taken before injecting the Gd-contrast agent. Frames 2-27 were taken with an interval of 4 min 36 s. The time courses of Gd-signals are shown as mean ± SEM (mean: dotted line, P4-weeks; solid line, P8-12-weeks, ± SEM, delineated shadow). The time points with a significant difference between the P4- and P8-12-weeks are asterisked (*p < 0.05). **B,** Summary bar graphs of accumulated Gd-signals for 36.8 minutes (from Frame 2 to Frame 9) at two developmental stages of each region (mean ± SEM, blank and filled bars for P4- and P8-12-weeks, respectively). Significant differences of “Parenchyma” were found in the Cerebellum mid (ranksum = 84, *p = 0.035), Spinal Cord (ranksum = 85, *p = 0.0426), SC (ranksum = 80, *p = 0.0149), VTA Right (ranksum = 83, *p = 0.0286), VTA Left (ranksum = 85, *p = 0.0426), Orbitofrontal Right (ranksum = 77, *p = 0.0072), Orbitofrontal Left (ranksum = 80, *p = 0.0149), mPFC Left (ranksum = 85, *p = 0.0426), Olfactory bulb Right (ranksum = 72, *p = 0.0019), Olfactory bulb Left (ranksum = 73, *p = 0.0025), and Olfactory bulb center (ranksum = 72, *p = 0.0019). Otherwise, not significant.

**Figure 5.**
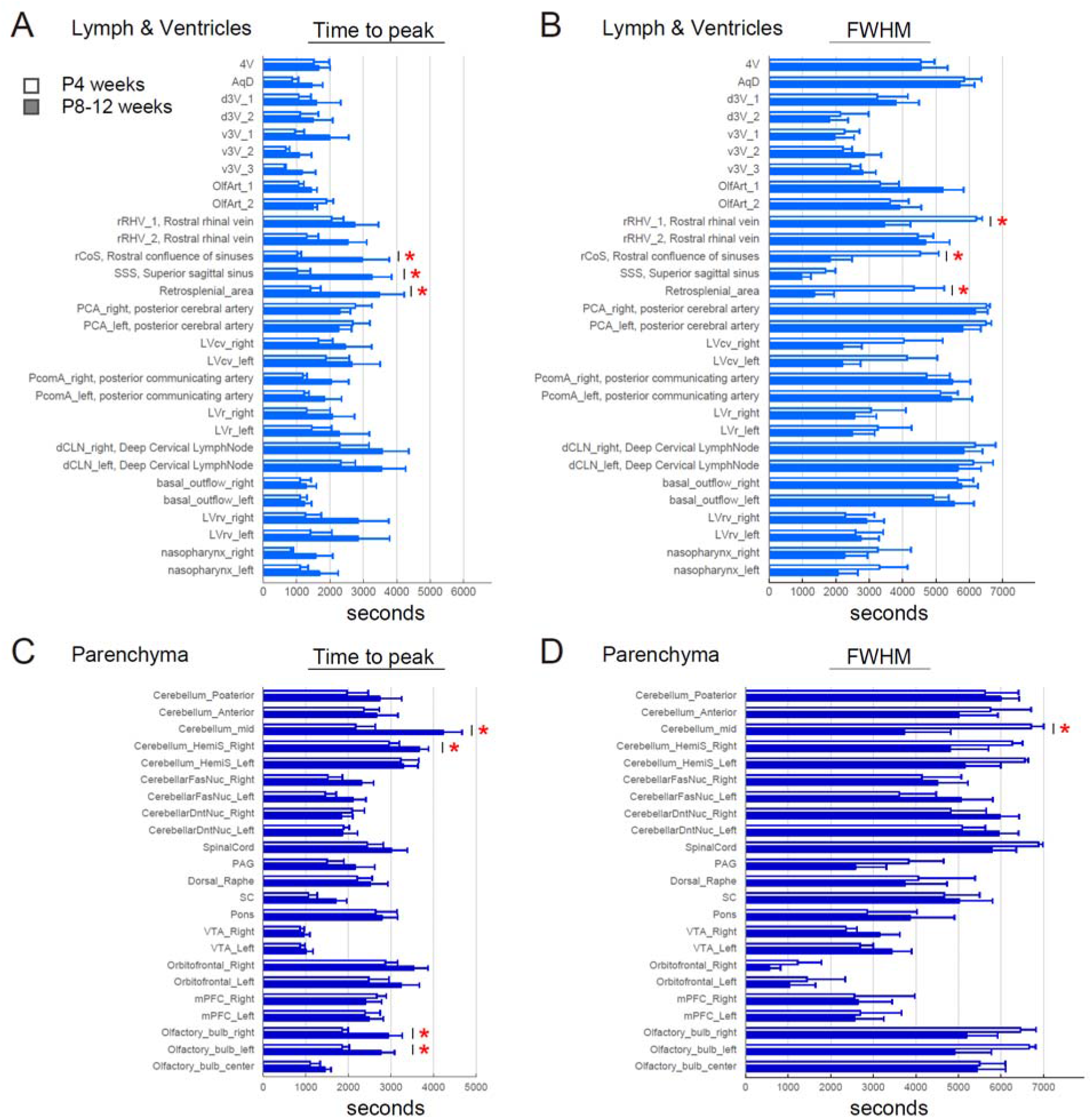
Time to peak and full width at half maximum (FWHM) of all the normalized Gd signals at different brain regions of the lymphatic vessels and ventricles, and parenchymas at two developmental stages. All the times to peak and FWHMs are shown in each group by developmental stages (P4-vs P8-12-weeks) and by vasculature – parenchyma axis for the comparison of the waveforms in each brain region. Bar graphs and error bars show mean ± SEM of time to peak (A and C) and FWHM (B and D) (Blank bar: P4-weeks. Filled bar: P8-12-weeks.). Asterisks indicate significant differences at the level of p < 0.05 by two-tailed Student’s *t*-test with unequal variances because two sampled groups are from the same distribution. In A of the time to peak, significant differences of “Lymph & Ventricles” were found in the rCoS (t(10.4) = 2.331, *p = 0.0411), SSS (t(12.7) = 3.089, *p = 0.0088), and retrosplenial area (t(11.8) = 2.564, *p = 0.0252). Otherwise, not significant. In B of the FWHM, significant differences of “Lymph & Ventricles” were found in the rRHV_1 (t(11.1) = −3.393, *p = 0.0059), rCoS (t(14.0) = −3.085, *p = 0.0081), and retrosplenial area (t(11.0) = −2.746, *p = 0.0190). Otherwise, not significant. In C of the time to peak, significant differences of “Parenchyma” were found in the Cerebellum_mid (t(14.1) = 3.180, *p = 0.0066), Cerebellum_HemiS_Right (t(14.3) = 2.173, *p = 0.0470), Olfactory_bulb_right (t(13.3) = 2.996, *p = 0.0101), and Olfactory_bulb_leftt (t(14.4) = 2.497, *p = 0.0252). Otherwise, not significant. In D of the FWHM, significant differences of “Parenchyma” were found in the Cerebellum_mid (t(8.0) = −2.652, *p = 0.0291). Otherwise, not significant.

**Extended Data Figure 4-1.**
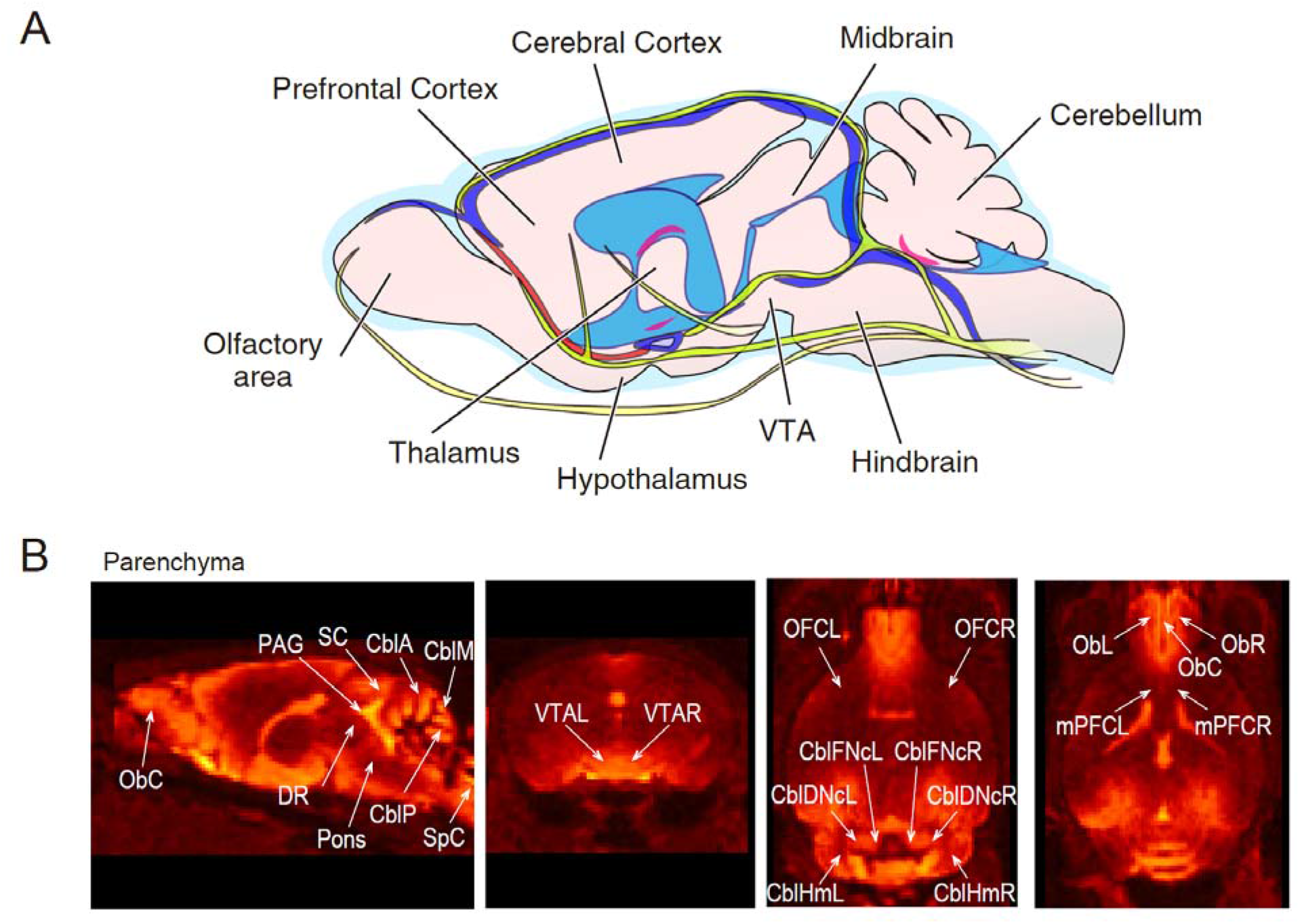
Schematic illustration of brain parenchymal regions and illustrations of brain parenchymas with interest, Related to Figure 4. CblA, anterior lobes of the cerebellar cortex. CblM, middle region (superior region of the posterior lobes) of the cerebellar cortex. CblP, posterior lobes of the cerebellar cortex. CblHm, cerebellar hemisphere. CblDN, dentate nucleus. CblFNc, fastigial nucleus. Dorsal Raphe, dorsal raphe nucleus. mPFC, medial prefrontal cortex. Ob, olfactory bulb. OFC, orbitofrontal cortex. PAG, periaqueductal gray. SC, superior colliculus. SpC, spinal cord. VTA, ventral tegmental area. R, C, and L indicate the right, center, and left, respectively.

Cerebellum Posterior, posterior lobes of the cerebellar cortex. Cerebellum Anterior, anterior lobes of the cerebellar cortex. Cerebellum mid, middle region (superior region of the posterior lobes) of the cerebellar cortex. Cerebellum HemiS, cerebellar hemisphere. CerebellarFasNuc, fastigial nucleus. CerebellarDntNuc, dentate nucleus. SpinalCord, spinal cord. PAG, periaqueductal gray. Dorsal Raphe, dorsal raphe nucleus. SC, superior colliculus. VTA, ventral tegmental area. Orbitofrontal, orbitofrontal cortex. mPFC, medial prefrontal cortex.

**Extended Data Figure 5-1.**
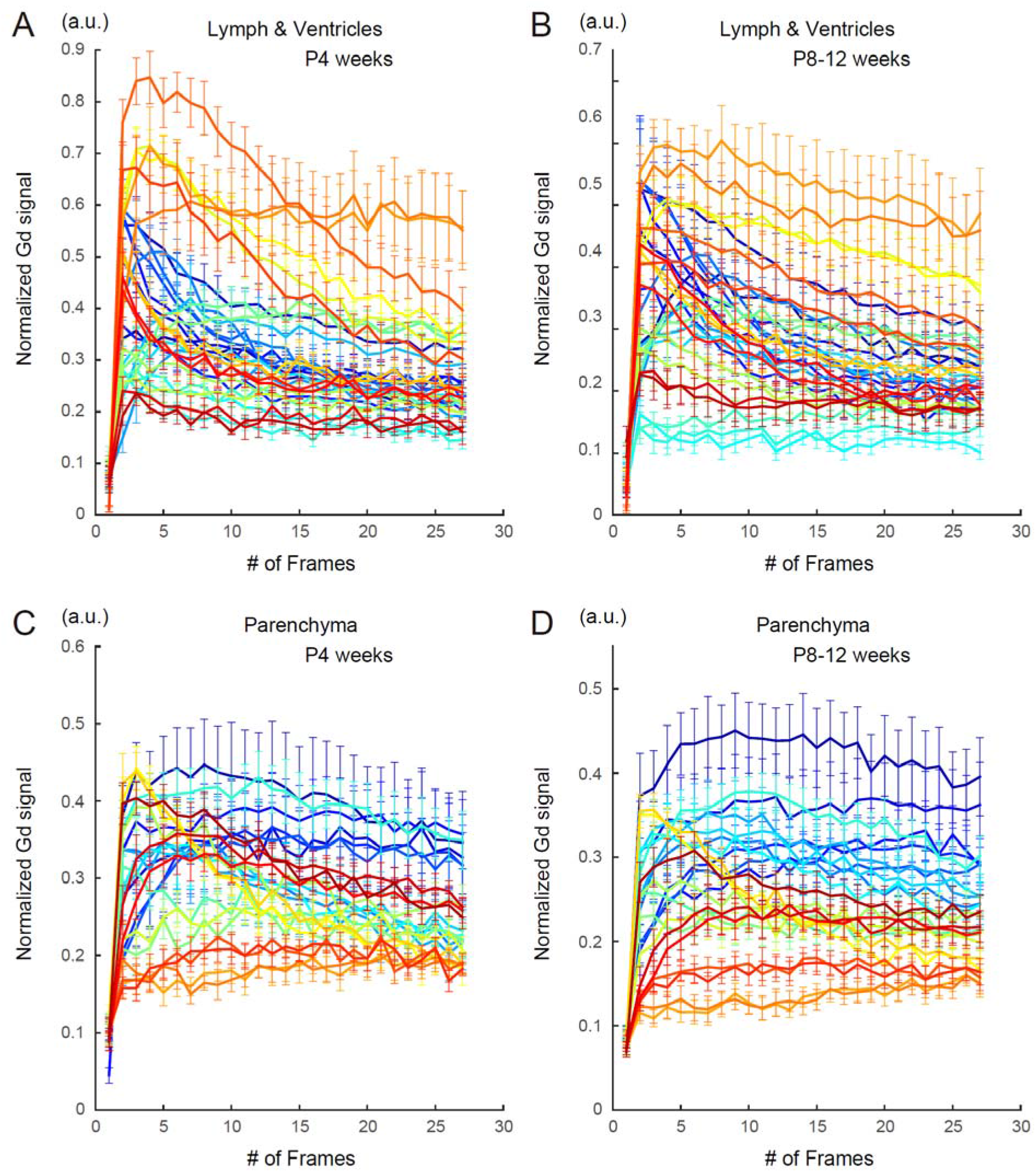
Time-courses of all the normalized Gd signals at different brain regions of the lymphatic vessels and ventricles and parenchymas at two developmental stages (P4 and P8-12-weeks), Related to Figure 5. All the time courses shown in Figures 3A and 4A are classified and displayed in each group by developmental stages and by vasculature – parenchyma axis for the comparison of the waveforms in each brain region. The color code is matched to that in Figures 3 & 4.

Next, to examine the time courses of the Gd-signals in the brain parenchyma, we took 23 points of interest (**Figure 4**; **Extended Data Figure 4-1**) and measured the normalized Gd-signals of the cerebellar cortex, cerebellar nuclei, spinal cord, periaqueductal gray, dorsal raphe, superior colliculus (SC), Pons, ventral tegmental area (VTA), orbitofrontal cortex, medial prefrontal cortex (mPFC), and olfactory bulb (as “Parenchyma”). There is also a wide range of variance in the normalized signal intensity of the regions of Parenchyma. The normalized Gd-signals were also high in the basal brain regions, such as VTA. On the other hand, the normalized Gd-signals were also high intensity in the posterior brain regions and low in the frontal cortex, reflecting that the Gd-signals in the parenchyma were polarized in the rostral-caudal axis in the cisterna magna injections (Kress et al., 2014). The normalized Gd-signals were significantly higher in P4-week-old animals than that in P8-12-week-old animals in the regions of SC, VTA, orbitofrontal cortex, and olfactory bulbs. In the middle lobules of the cerebellar cortex (the superior region of the posterior lobes), spinal cord, pons, and mPFC showed transient but significantly higher Gd-signals in the young animals (**Figure 4A**). The time courses of Gd-signals appeared faster in the brain vasculatures (in lymphatic vessels & ventricles, **Figure 3A**) than those of the brain parenchymal area expected for the VTAs (**Figure 4A**; **Figure 5**), suggesting slow infiltration of the CSF into the parenchyma. Compared to the other regions of parenchyma, the time courses of the cerebellar cortex and nuclei showed longer rise time to the peak and longer decay (Cerebellum mid, Cerebellum HemiS, CerebellarFasNuc, and CerebellarDntNuc in **Figure 4A** and **Figure 5C&D**). In the BBB, AQP-4 expressing at the endfeet of astrocytes dominantly filters the CSF at most of the brain parenchymal regions (Nielsen et al., 1997). A chicken study suggest a fewer level of AQP-4 expression in the cerebellar molecular layer but a high expression pattern along the entire pial surface, Bergmann glial fiber terminals, and perivascular processes facing the capillaries, which may occur in the mouse brain and may delay the infiltration (Nico et al., 2002). In Figure 4B and Figure 6, the bar graphs of the cumulative normalized Gd-signals of the initial and late period (from Frame 17 to Frame 27) again indicate the distinct regions with high Gd-signals in the young period: in the cerebellar cortex, spinal cord, SC, VTA, orbitofrontal cortex, mPFC, and olfactory bulb. Time to peak and FWHM of Gd-signals also support the parenchymal regions with high Gd-permeability (**Figure 5**; **Extended Data Figure 5-1**). As far as we know, no developmental differences in the cerebrovascular function and the vascular running have been reported during this experimental period. These results of the DCE-MRI imply that the immature barrier function of the vasculature system including the lymphatic vessels and the BBB allows the infiltration of low molecular weight substances and, perhaps, cells to infiltrate parenchyma in those areas.

**Figure 6.**
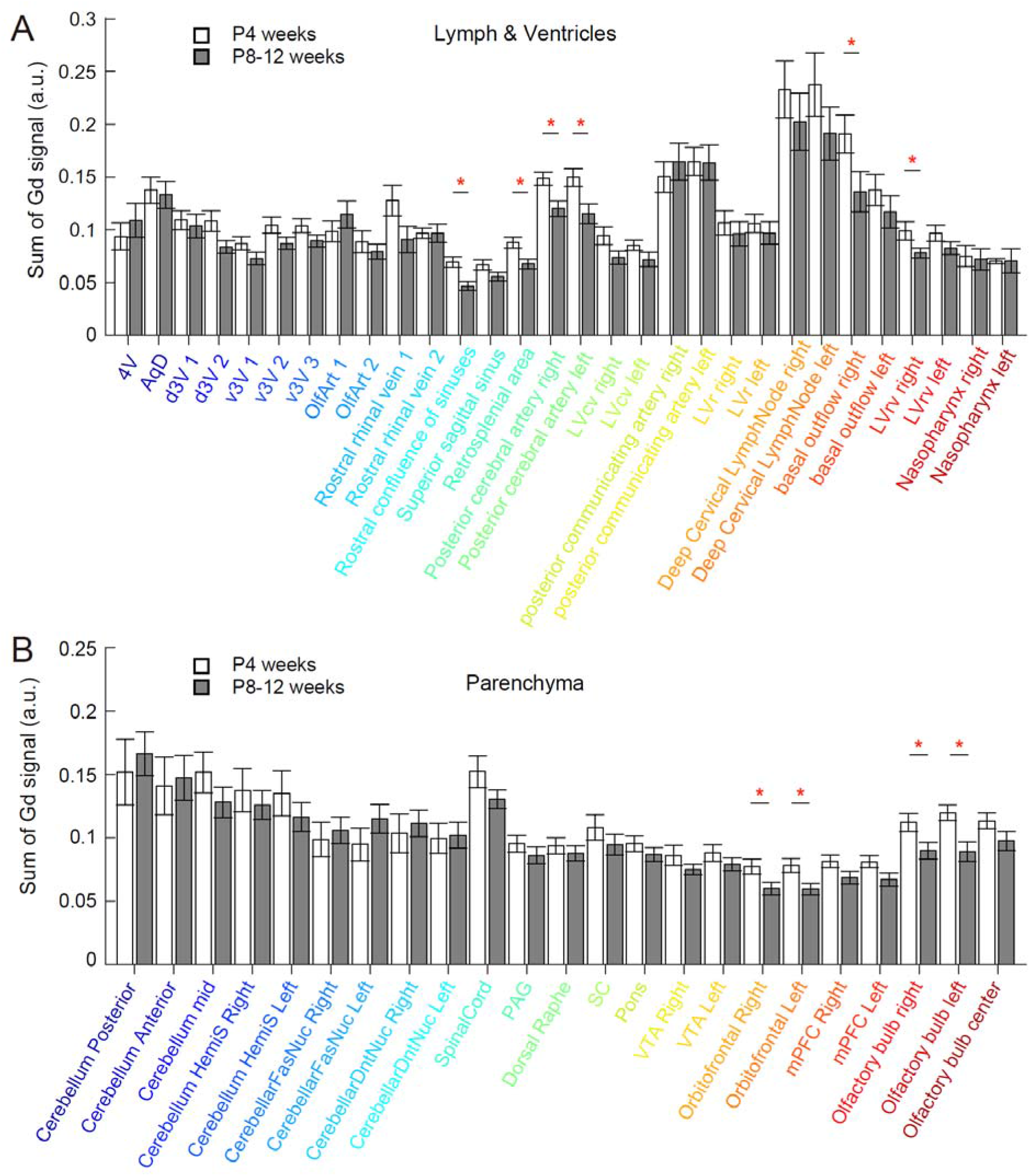
Summary bar graphs of accumulated Gd-signals in the late phase of experiments. **A,** Summary bar graphs of accumulated Gd-signals in the late phase of experiments for 50.6 minutes (from Frame 17 to Frame 27; i.e., from 77’00” to 124’00”) at two developmental stages of the lymphatic vessels and ventricles (mean ± SEM, blank and filled bars for P4- and P8-12-weeks, respectively). Significant differences of “Lymph & Ventricles” were found in the Rostral CoS (ranksum = 75, *p = 0.0043), retrosplenial area (ranksum = 80, *p = 0.0149), PcomA Right (ranksum = 80, *p = 0.0149), PcomA Left (ranksum = 81, *p = 0.0186), LVrv right (ranksum = 84, *p = 0.035), basal outflow Right (ranksum = 85, *p = 0.0426), and LVrv right (ranksum = 83, *p = 0.0286). Otherwise, not significant. **B,** Summary bar graphs of accumulated Gd-signals in the late phase of experiments of the parenchymas. Significant differences of “Parenchyma” were found in the Orbitofrontal Right (ranksum = 83, *p = 0.0286), Orbitofrontal Left (ranksum = 79, *p = 0.0118), Olfactory bulb Right (ranksum = 80, *p = 0.0149), and Olfactory bulb Left (ranksum = 78, *p = 0.0093). Otherwise, not significant.

Lastly, we confirmed no microhemorrhages in immature brains in vivo by visualizing the artery distribution in P4-week mouse brains using MRA. Since MRA contrasts the fast blood fluid flow, it mainly indicates the arterial blood flow. Our observation did not find any leakage of the blood vessels (**Figure 7**), suggesting that the artery is intact and other perivascular space such as capillary, the subarachnoid space, and meningeal lymphatic vessels would be the sites of leakage.

**Figure 7.**
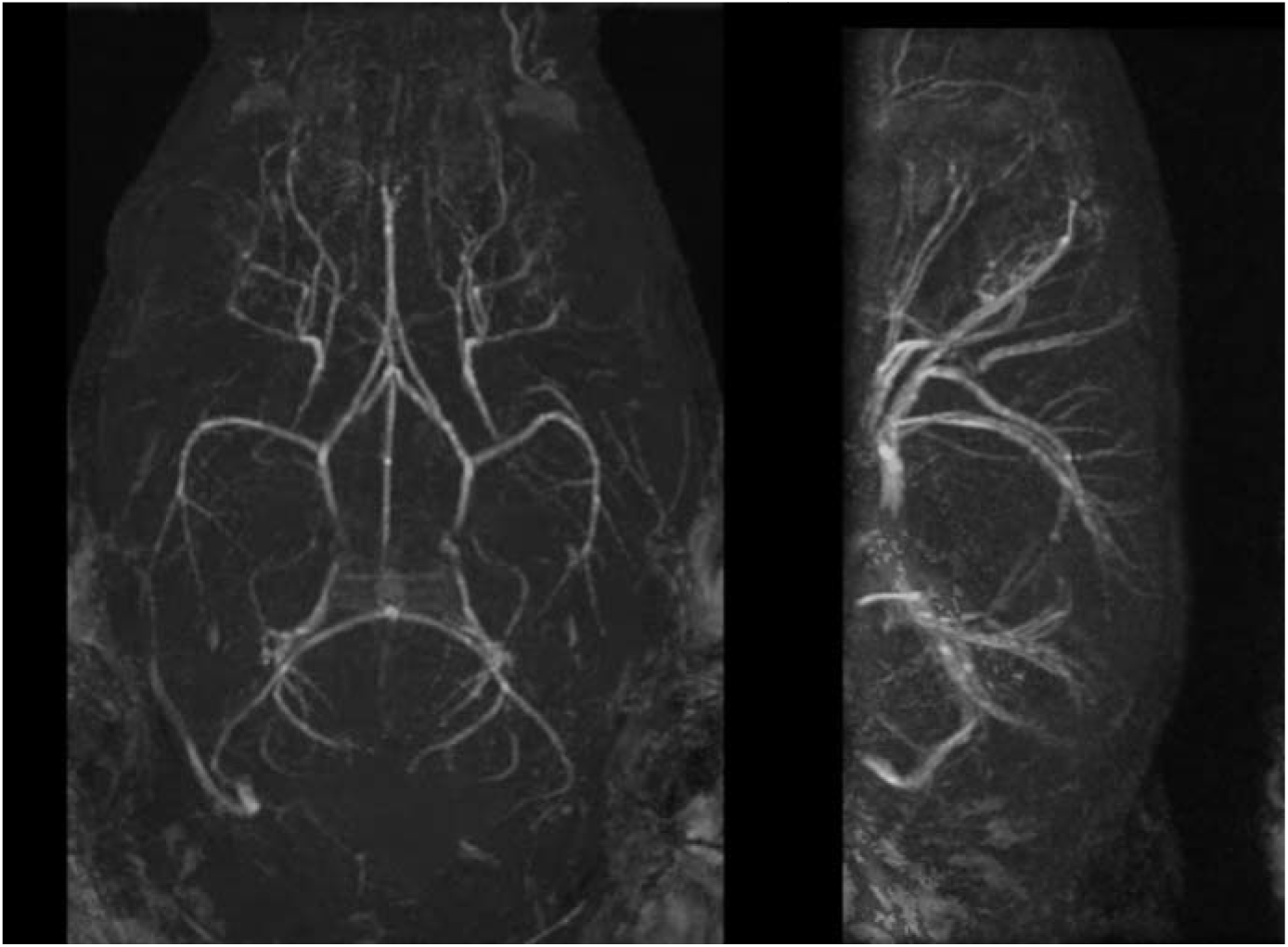
MR angiograms of a young (P4-weeks) C57BL/6 mouse. Horizontal (*left*) and sagittal (*right*) views of representative MRA projection images are displayed. No detectable microhemorrhages were observed in the brain arteries. Isotropic spatial resolution of 100 μm.

## Discussion

In this study, we monitored the bulk CSF flow using DCE-MRI and compared the Gd-signals along with the brain lymphatic flow and in the brain parenchymal regions. According to Stanton et al. (2021), the Gd-contrast agent injected into the cisterna magna is distributed along with the perivascular space of the blood vessels in the subarachnoid space cisterns, and the contrast agent infiltrates into the parenchyma. In P4-week animals, the Gd-signals showed higher intensity in distinct regions of brain parenchyma, such as the olfactory bulb, prefrontal cortex, parietal surface regions of the neocortex along with SSS, ventral midbrain, and a part of cerebellum, as well as in the basal brain regions, compared to mature P8-to P12-week-old mice (**Figures 3–6**). No substantial leakage of the arterial flow by MRA (**Figure 7**) implied that the infiltration of Gd-signals comes from the subarachnoid space, perivascular space along blood vessels and capillaries, and meningeal lymphatic vessels. Our results suggest that the CSF barrier in mouse brains is still permeable until P4 weeks.

### A distinct leak in Gd-signals even at the timing when the BBB integrity is established

Previous studies by Bauer et al. (1995) using the injection of Trypan blue via the umbilical vein suggested that the staining substance does not infiltrate into the CSF space from the systemic circulation, and such a barrier is established in the early embryonic stage. Although injected Trypan blue had been excluded from most parts of the central nervous system (CNS), those experiments did not necessarily indicate the impermeability of CSF into the brain parenchyma nor the functional completion of BBB. On the other hand, electrical resistance across BBB in anesthetized rodents was also measured previously (Butt et al., 1990), and the averaged transendothelial electrical resistance (TEER) at one-month-old was higher up to five times than that immediately before the birth. The result had been regarded to demonstrate the rapid maturation of BBB. However, it did neither necessarily indicate the completion of all the BBB-barriers because measured resistance showed a great variance among the samples (30-5900 Ωcm^2^). Most of the TEER of one-month-old rats was less than 1000 Ωcm^2^, and the TEER of veins (800 Ωcm^2^) was significantly lower than that of the artery (Butt et al., 1990). The barrier should become functional during postnatal maturation, but it had been unclear at which developmental stage the BBB-barrier is fully established (Bauer HC et al., 2014). In this study, DCE-MRI allowed us to visualize how the small molecule spreads and infiltrates into the CSF space in the entire brain region. And we found distinct Gd-signals in parenchymas at P4 weeks, which is the later stage than the timing when normally believed the BBB is completed. However, a low resolution of the MRI method limits our interpretation of how the Gd-contrast agent permeates into the parenchyma and with what machinery works for the developmental changes.

### A possibility of the permeability along the immature meningeal lymphatic vessels

Due to the functional identities, there are two canonical vasculature systems in the brain: the meningeal lymphatic vessels and the glymphatic system (Louveau et al., 2017). Given a certain leak across to the parenchymal regions as the underlying mechanism, we may raise the following assumptions to account for our observations of the difference between P4-week and P8-12-week mouse brains. One possibility is that the meningeal lymphatic vessels are still permeable at postnatal development. In CSF production at the blood-CSF barrier, the choroid plexus absorbs water via aquaporin (AQP)-1 and ions from the blood, generating CSF in the ventricles and subarachnoid spaces (Dietz et al., 2020; Segawa et al., 2021). CSF circulates via thin lymphatic ducts in the meninges (: meningeal lymphatic vessels). The flow stream of CSF and meningeal lymph is thought substantially driven by the pulsatile and cerebral perfusion pressure. CSF in the subarachnoid space flows into the venous sinus via arachnoid granulations (Rustenhoven et al., 2021; Ringstad & Eide, 2020). The recent rediscovery of the meningeal lymph elucidated that the lymphatic network scavenges macromolecules and waste products of tissues and secures immune cells into the CNS (Louveau et al., 2015, 2018; Aspelund et al., 2015; Absinta et al., 2017; Da Mesquita et al., 2018b). The leaky barrier function of meningeal lymphatic vessels and the reduction of brain perfusion of CSF macromolecules in aged brains are suggested as a hallmark of the cognitive impairment (Da Mesquita et al., 2018a; Sweeney et al., 2018; Ahn et al., 2019). Lymphatic endothelial cells (LECs) form both button-like junctions (high permeability for the initial lymphatic vessels) and zipper-like (: tight barrier) junctions, both of which are composed of adherens and tight junctions. In mature P3-month-old mice, more than 60% of junctions between LECs of dorsal meningeal lymphatic vessels are zipper-like, and around 20% are button-like (Ahn et al., 2019), suggesting the leakage would be shut-down as seen in P2-to P3-month-old mice of our results. Such a button-like structure in the dorsal forebrain may allow infiltration of the Gd-contrast agent. Therefore, the meningeal lymph and associated veins in the juvenile brains may be leaky because of the dominance of button-like junctions or their immaturity.

### A possibility of functionally immature BBB

The glymphatic system is another brain perfusion machinery for CSF (Iliff et al., 2012, 2013). The Gd-contrast agent flows along with the perivascular space of the blood vessels in the subarachnoid space cisterns and possibly infiltrates into parenchyma via the BBB. Thus, the second possibility is that the leaky CSF barrier between the perivascular space and parenchyma. Vasculatures and capillaries are coated by the astrocytic endfeet, and the AQP-4 existing there is the major gateway for the CSF/ISF exchange of the BBB. AQP-4 expresses in the embryonic mouse brain at E16, and a polarized localization for organizing the water gateway emerges at P1-to P3-days (Fallier-Becker et al., 2014). A study by Kress et al. (2014) suggested the attenuated CSF tracer influx into the parenchyma in the course of development from 2-to 3-month-to 18-month-old mice due to loss of perivascular AQP-4 polarization in the aged brains. Contrary to this, our results indicate higher permeability in one-month-old mice in distinct brain regions.

The BBB is tightly regulated by the expansion of the microvascular network and the volume expansion of the brain during maturation (Coelho-Santos & Shih, 2020). Over weeks after birth, the cortical microvasculature network undergoes dramatic expansion via capillary angiogenesis (Risau & Wolburg, 1990; Risau, 1997; Marín-Padilla, 2012; Harb et al., 2013). Volumetric measurements suggested that the mouse brain keeps expanding at least until the end of postnatal one month, and the rate is high in the neocortex and the cerebellum (Zhang et al., 2005). Thus, maturation of the cerebrovascular capillaries lasts during the postnatal period (Coelho-Santos & Shih, 2020). Considering the infiltration of the CSF from the perivascular space to the parenchymas, the functionality of the BBB could be incomplete during that period. Our findings of the high intensity of Gd-signals in the juvenile brain in the olfactory bulbs, mPFC, orbitofrontal cortex, the upper part of the cingulate cortex nearby the longitudinal fissure (i.e., rostral CoS, SSS, and retrosplenial area), SC, VTA, peripheral regions of the posterior cerebral artery (: the border regions between the midbrain and posterior hippocampus), and middle lobules of the cerebellum suggest the leaky barrier in those regions (**Figure 4**). Although our MRA observation did not support any microvasculature leakage in the arteries (**Figure 7**), the CSF infiltration may occur in the microstructure at a slow speed.

Strand breaks of tight junctions (TJs) are known to occur during maturation and inflammatory states of diseases (Huber et al., 2001; Varadarajan et al., 2019; Greene et al., 2019). TJs between cerebrovascular endothelial cells as blood endothelial cells (BECs) form a strong barrier (Butt et al., 1990). Besides, the interaction of astrocyte endfeet and pericytes to BECs leads to the formation of BBB with a fenestrated barrier. The junctions between BECs include TJs, adherens junctions, and gap junctions. The TJs are composed of claudins, occludin, and ZO-1 accompanied with JAMs. In BBB, a barrier-forming type claudin-5 is the major component (Nitta et al., 2003), while other claudins are also known to compose the BEC TJs (Berndt et al., 2019). In claudin-5 knockout and knockdown mice, the cerebrovascular permeability is significantly increased, and the BBB was revealed to permeate substances with at least up to 800 Da of the molecular weight (Nitta et al., 2003; Campbell et al., 2008). However, we have no ideas if such leakage or exchange of CSF and plasma component occurs via strand breaks and affects the infiltration of Gd-contrast agent in the intact developing brain.

### Relevance to brain physiological dysfunctions

Currently, researchers assume that disruption of BBB integrity, infiltration of immune cells, and inflammatory cytokines via aberrant immune activity in the developing brain have a link to various types of psychiatric disorders (Claudio, 1996; Perry et al., 1997; Lou et al., 1997; Bolton et al., 1998; Davalos et al., 2012; Lepennetier et al., 2019; Montagne et al., 2020; Yang et al., 2020; Mastorakos et al., 2021). Invasion of stimulated immune cells or inflammatory cytokines into the brain parenchyma may disturb the neuronal function through excessive immune responses (Yirmiya et al., 2015; Becher et al., 2017; Dietz et al., 2020; Segawa et al., 2021). The BBB TJ-disruption could be a signature phenotype of psychiatric disorders in animals. A loss of claudin-5 in the nucleus accumbens is shown associated with depressive-like behaviors, while loss of claudin-5 in the hippocampus and mPFC is associated with schizophrenia-like behaviors (Menard et al., 2017; Greene et al., 2018). Therefore, it was important to know the functional development of the barrier in the brain to display which regions are permeable. Aberrant immunity in the vulnerable areas would also lead to neurophysiological dysfunctions and disruption of functional connectivity. For instance, activated microglia or released inflammatory cytokines in the prefrontal cortex and cerebellum induce forms of plasticity of intrinsic excitability of various types of neurons (Schonewille et al., 2010; Belmeguenai et al., 2010; Ohtsuki et al., 2012; Duan et al., 2018; Yamamoto et al., 2019; Ohtsuki, 2020; Ohtsuki et al., 2020; Ozaki et al., 2021; Yamawaki et al., *in revision*), as well as of excitatory and inhibitory synaptic transmission (Zhang et al., 2014; Duan et al., 2018; Yamamoto et al., 2019; Zheng et al., 2021; Yamawaki et al., *in revision*). These modulations of both brain-wide functional networks are associated with psychiatric diseases-like behaviors (Yamamoto et al., 2019; Granja et al., 2021). One of the upcoming questions is if dysfunction of the vasculature system suffices to the emergence of developmental disorders via peripheral immunity and if the timing of the maturation of CSF barrier determines the period for the emergence of developmental disorders. Future comprehensive work would explain the relevance between CSF-flow dynamics in developmental brains and brain dysfunctions.

## Materials & Methods

### Animal ethics statement

All procedures were performed following the guidelines of the Animal Care and Use Committees and approved by the Ethical Committee of Kyoto University. All animal handling and reporting comply with the ARRIVE guidelines. Mice were housed (4 animals at maximum in each cage) and maintained under a 12-h light: 12-h dark cycle, at a constant temperature and humidity (20−24 □, 35%−55%), with food and water available ad libitum. We conducted all animal procedures for MRI experiments following the guidelines of animal experiments at Kyoto University. The MRI experiments of this work were performed in the Division for Small Animal MRI, Medical Research Support Center, Graduate School of Medicine, Kyoto University, Japan.

### Gd-reagent injection

For injection of Gd-contrast agent into the cisterna magna, C57BL/6 male mice were fixed to the stereotaxic apparatus (NARISHIGE Group, Japan) under inhalation isoflurane (2% for induction: 0.8% for operation with 0.1–0.2% oxygen or air). After cutting off the scalp and lesser posterior rectus capitis muscles of the back of the mouse with scissors and a surgical knife, we made one tiny sting (with a 200 μm diameter) by penetration using a 27-gade needle, following which the tip of the needle of a syringe (GASTIGHT^®^ #1705, Hamilton Co.) was inserted 3.6–3.9 mm forward at a slow angle of 5° negative. We infused a 35–38 μl undiluted Gd-contrast agent (GadoSpin M, Miltenyi Biotec, USA) in 5–8 minutes. Less amount of injection did not achieve the lateral ventricles. To monitor the entire CSF flow dynamics in the entire brain vasculatures, we intentionally infused a bulk amount of Gd-contrast agent. Please note that this rate and duration of Gd-contrast agent infusion would lead to reflux of subarachnoid CSF back into the ventricular CSF compartments, indicating that the physiological direction of CSF-flow is not maintained initially (Kress et al., 2014). The wound was sealed with Spongel (Astellas Pharma Inc.), after which the animals were allowed to proceed with the MR imaging. The maximum time interval for the start of the initial MRI acquisition from the end of the operation was 9 minutes, and thus, we used this time in Figure 2. Upon intravenous injection, GadoSpin M is rapidly distributed in the extracellular space. According to the prescription of GadoSpin M, the Gd-contrast agent is excreted via glomerular filtration (kidneys) within hours. We did not find any plasma protein complex during the MRI.

### MRI experiments

#### 1. Animal preparations

Under general anesthesia with inhalation of 2.5 % isoflurane in the air at 1 L/min through a face mask, mice were placed in a cradle in a prone position. The mouse head was fixed using a tooth bar and surgical tapes. The animal’s respiratory rate and rectal temperature were continuously monitored via a pressure-sensitive respiration sensor and thermistor temperature probe, respectively, and monitored using a dedicated system (Model 1025, MR-compatible Small Animal Monitoring & Gating System; SA Instruments, Inc., NY, USA) equipped with software (PC-SAM V.5.12; SA Instruments). The body temperature was maintained by a flow of warm air using a heater system (MR-compatible Small Animal Heating System, SA Instruments). The isoflurane concentration was adjusted in the range of 0.6 − 1.2 % to ensure a stable and reproducible depth of anesthesia based on the respiration rate.

#### 2. MRI acquisitions

All MRI measurements were conducted on a 7-Tesla preclinical scanner (BioSpec 70/20 USR; Bruker BioSpin MRI GmbH, Ettlingen, Germany). quadrature transmit-receive volume coil (inner diameter 35 mm, T9988; Bruker BioSpin) was used to detect MR signals. The MRI system was controlled with a dedicated operation software (ParaVision 5.1; Bruker BioSpin).

##### 2.1. 3D T1-weighted MRI

A series of 3D T1-weighted (T1W) images was acquired using a fast low angle shot (FLASH) sequence with the following acquisition parameters: repetition time (TR), 30 ms; echo time (TE), 3.3ms; flip angle, 25°; field of view (FOV), 22.5 × 15 ×15 mm^3^, acquisition matrix size, 144 × 96 × 96; isotropic spatial resolution of 0.15625 mm; coronal orientation (i.e., axial in mouse brain); without averaging; and scan time, 4 min 36 s. As a baseline image as Frame 1, a non-contrast-enhanced image was also acquired before injecting the Gd-contrast agent. The FLASH scan was repeated 26 times over 120 min with 4 min 36 s intervals as Frames 2 to 27.

##### 2.2. 3D T2-weighted MRI

As an anatomical reference image, a 3D T2-weighted (T2W) image was acquired using a rapid acquisition with relaxation enhancement (RARE) sequence. The acquisition parameters were: TR, 1500 ms; effective TE, 45 ms, RARE factor, 16; fat suppression; flip back; and scan time, 14 min 24 s. The other parameters were the same as the 3D T1W imaging.

#### 3. Image processing

Using the acquired MR images, the following analyses were conducted. All data were processed using FMRIB Software Library (FSL; the Analysis Group, FMRIB, Oxford, UK, https://fsl.fmrib.ox.ac.uk/fsl/fslwiki) (Jenkinson et al., 2012) and ImageJ (Rasband, W.S., ImageJ, U. S. National Institutes of Health, Bethesda, Maryland, USA, https://imagej.nih.gov/ij/, 1997-2018).

##### 3.1. Pre-processing for analyzing the time course of contrast enhancement

In the analyses using FSL, all images had their pixel dimensions scaled up in the NIfTI header by a factor of 10 to avoid the scale-depended issue when using standard FSL software. Pixel values of the baseline image and the series of contrast-enhanced images were converted to the absolute image intensity. The baseline image was linearly (6 degrees of freedom) registered to the first image in the contrast-enhanced time series data set using FSL’s FLIRT (Jenkinson et al., 2002). These images were motion-corrected using FSL’s MCFLIRT (Jenkinson et al., 2002) and then a mean T1W image was generated by taking the mean value across time on a pixel-by-pixel basis. The mean T1W image was linearly (6 degrees of freedom) registered to the corresponding anatomical (T2W) image and the transformation matrix was stored. The transformation matrix was used for the registration of the time series T1W images to the anatomical image. A T2W brain template image of C57Bl/6 mouse (Hikishima et al., 2017), which is available from the Neuroimaging Informatics Tools and Resources Clearinghouse (NITRC) website (https://www.nitrc.org/projects/tpmmouse), was used in the present study. The template image was downsampled so as the spatial resolution to match our images. Then a rectangular region of interest (ROI), in which almost the minimum region surrounding the whole brain was included, was extracted from the template image. The anatomical image of each mouse was linearly (12 degrees of freedom) registered to the template brain and the generated transformation matrix was applied to the motion-corrected and registered T1W data series. All the processing for image registration was performed by using FSL’s FLIRT.

##### 3.2. Maximum intensity projection

To visualize the distribution of the Gd-contrast agent, maximum intensity projection (MIP) images were generated from a 3D T1W time series data set. Using 3D projection in ImageJ’s routine, a total of 36 MIP images for each volume (i.e., each time point data) was created through a total rotation angle of 360° with an increment of 10° and an axis of rotation along the anterior-posterior direction. The resulting data set depicts 360°-rotation of the MIP image.

##### 3.3. Data analyses and statistics

After preprocessing with FLIRT, we obtained the Gd-signals of the regions of interest with the FSLeyes (https://fsl.fmrib.ox.ac.uk/fsl/fslwiki/FSLeyes). The time series of the voxel intensities were taken from 4D NIFTI images of the lymphatic vessels, ventricles, and parenchymas. The coordination of the C57Bl/6 mouse brain template space (Hikishima et al., 2017) is followings (*i.e*., x, y, and z):

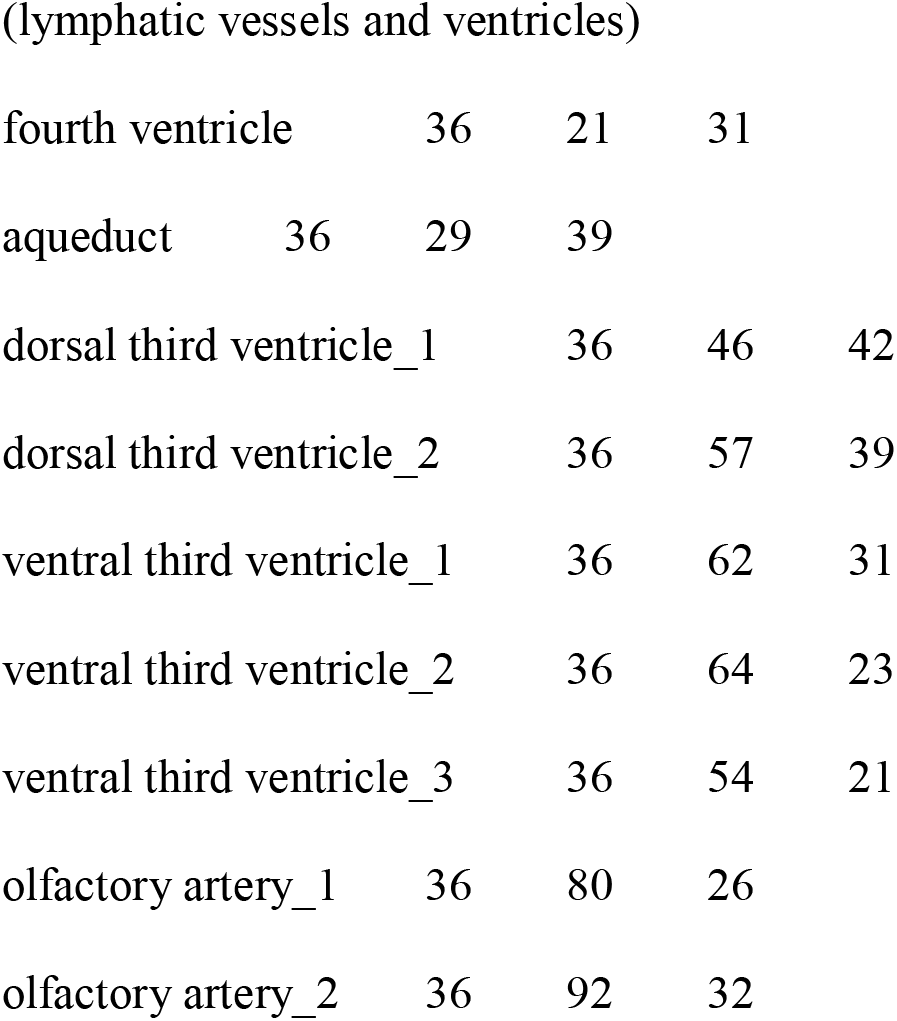
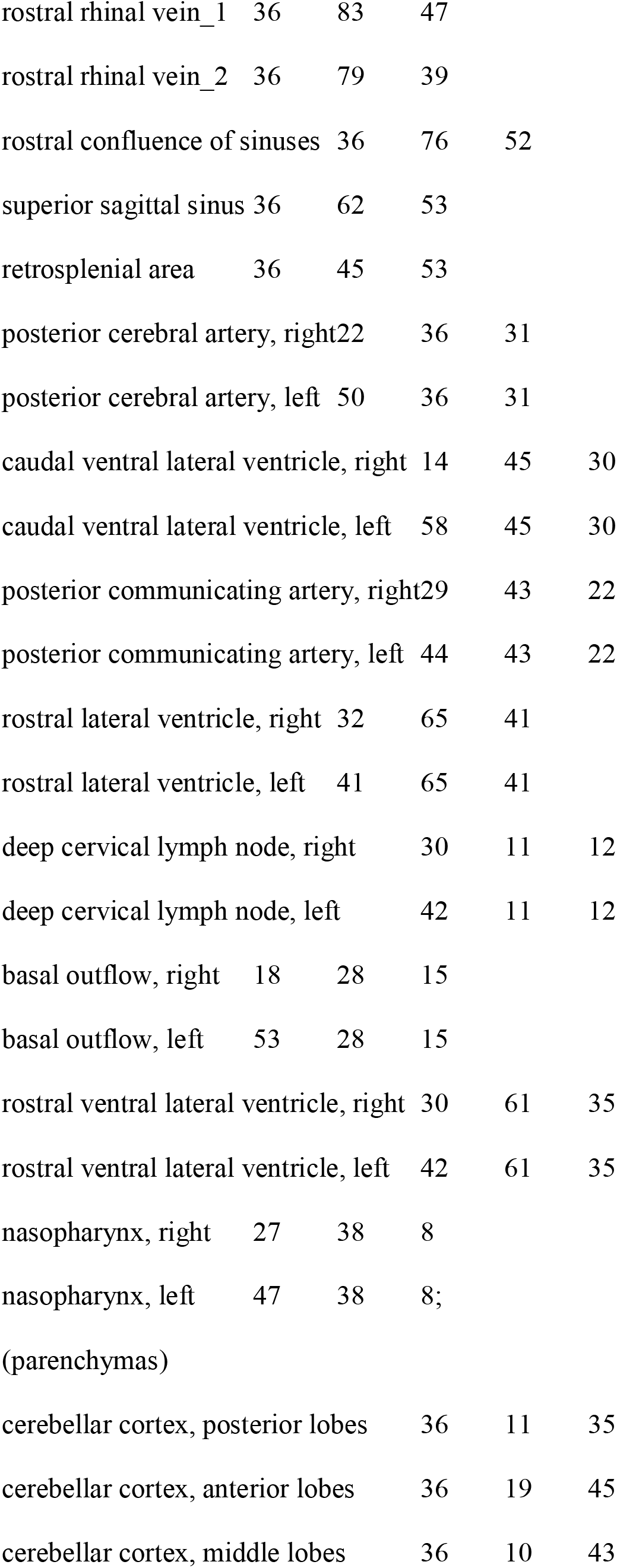
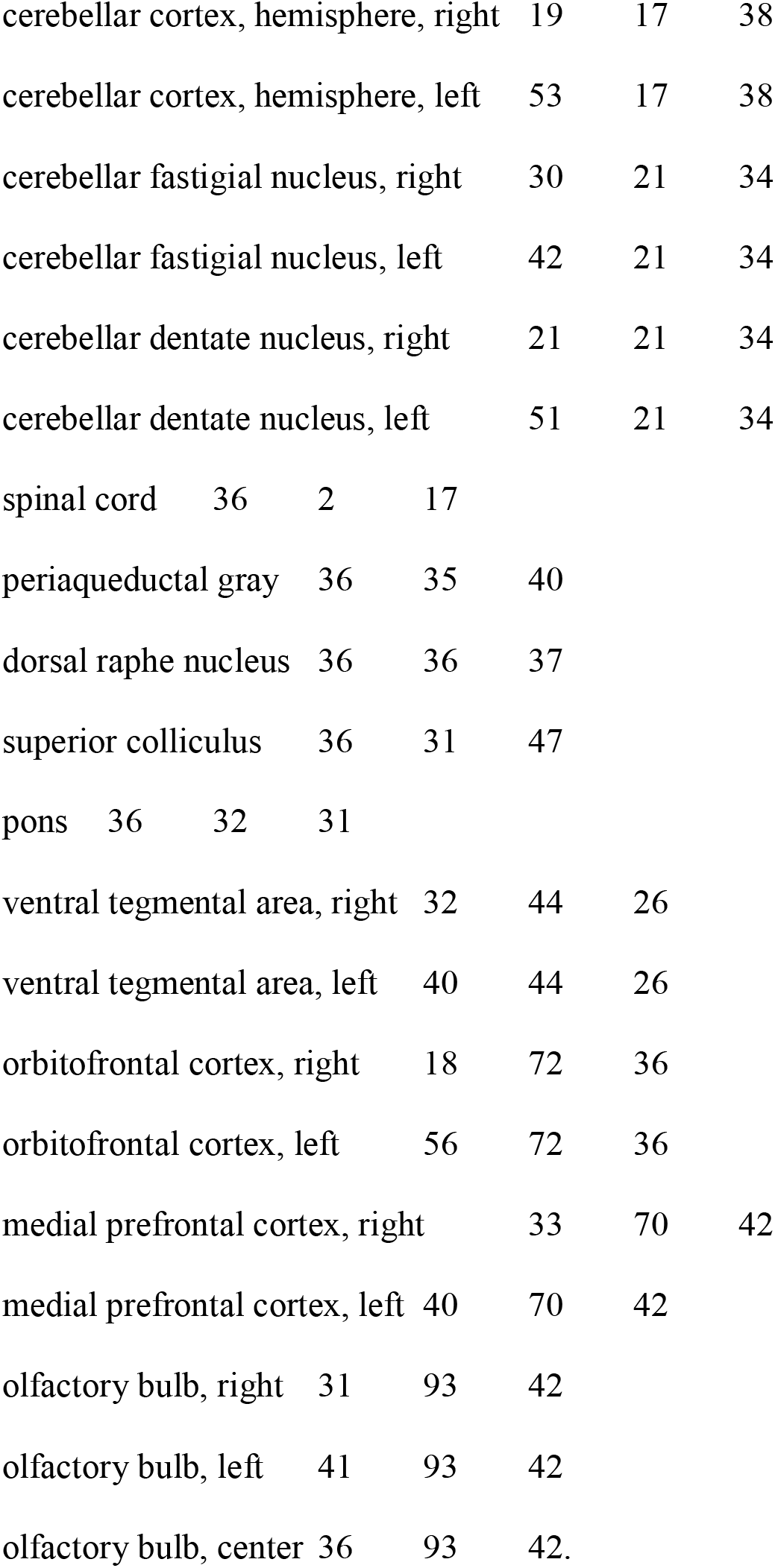

Due to the difference in the position and shape of lymphatic nodes, we adjusted the coordination to the center of the deep cervical lymph node of each animal. Voxel intensities as Gd-signals were normalized by both minimum and maximum intensity for each animal. Then, the average across animals and standard error of measurement (SEM) were taken for each brain region. All data are presented as mean ± SEM. Data of P4-week, P8-9-week, and P12-week-old mice were collected from 7, 5, and 6 animals for each. We summed up the P8-9-week and P12-week-old mice data together. One-sided Mann-Whitney-tests (Cardillo, 2009) were used to compare data of normalized Gd-signal amplitude and summed Gd signals between two independent groups (**Figure 3, 4, and 6**). Two-tailed Student’s t-tests with unequal variances were used for the data of time to peak and full-width of half maximum (FWHM) because two-sampled groups are considered from the same distribution (**Figure 5**). We considered p < 0.05 as the significance level throughout the study. All statistical analyses were performed using Matlab 2015.

#### 4. MRA

Three-dimensional time-of-flight MR angiography (3D TOF-MRA) was performed with the following acquisition parameters: pulse sequence, the gradient-echo sequence with a first-order flow compensation in all logical gradient directions; TR, 100 ms; TE, 4.63 ms; flip angle, 45°; FOV, 19.2×12.8×9.6 mm^3^; acquisition matrix size, 192×128×96; isotropic spatial resolution of 100 μm; the number of averages, 4; and scan time, 1 h 22 min.

## Acknowledgments

We would like to present our deep appreciation to Prof. Shu Narumiya, Drs. Takeshi Sakurai, and Kosuke Tanigaki for invaluable comments and supports on the research. We thank Rie Toyoda, and Konomi Nimura for the validation of data analyses and image drawings. MRI was performed at the Medical Research Support Center, Graduate School of Medicine, Kyoto University, which was supported by Platform for Drug Discovery, Informatics, and Structural Life Science from the Ministry of Education, Culture, Sports, Science and Technology, Japan. We also appreciate Sumitomo Dainippon Pharma Co., Ltd., ONO PHARMACEUTICAL CO., LTD., Mitsubishi Tanabe Pharma Corporation., and KYORIN Pharmaceutical Co., Ltd. as the sponsors for the Department of Drug Discovery Medicine.

## Author Contributions

G.O., H.I., Y.Y., and S.M. conducted the experiments. G.O., Y.W., Md S.A.P., and Y.Y. analyzed data. G.O. and H.I. designed the experiments. G.O., Y.W., H.I., Y.Y., Md S.A.P., and S.M. wrote the manuscript. This work was supported by grants from the Naito Foundation, the Mitsubishi Foundation, and the Takeda Science Foundation (all to G.O.), and by JSPS WISE program “The Graduate Program for Medical Innovation” (to Y.Y.). The funders had no role in study design, decision to publish, or preparation of the manuscript.

---

**Video S1, Related to Figure 1B.**

A representative movie of the initial frame of Gd-contrast images of the postnatal 4 weeks mouse. The movies were made from rotated series T1-images (3D maximum intensity projection images) with every 20-degree difference.

**Video S2, Related to Figure 1C.**

A representative movie of the initial frame of Gd-contrast images of the postnatal 9 weeks mouse. The movies were made from rotated series T1-images (3D maximum intensity projection images) with every 20-degree difference.

